# Optimization of Non-Coding Regions Improves Protective Efficacy of an mRNA SARS-CoV-2 Vaccine in Nonhuman Primates

**DOI:** 10.1101/2021.08.13.456316

**Authors:** Makda S. Gebre, Susanne Rauch, Nicole Roth, Jingyou Yu, Abishek Chandrashekar, Noe B. Mercado, Xuan He, Jinyan Liu, Katherine McMahan, Amanda Martinot, Tori Giffin, David Hope, Shivani Patel, Daniel Sellers, Owen Sanborn, Julia Barrett, Xiaowen Liu, Andrew C. Cole, Laurent Pessaint, Daniel Valentin, Zack Flinchbaugh, Jake Yalley-Ogunro, Jeanne Muench, Renita Brown, Anthony Cook, Elyse Teow, Hanne Andersen, Mark G. Lewis, Stefan O. Mueller, Benjamin Petsch, Dan H. Barouch

## Abstract

The CVnCoV (CureVac) mRNA vaccine for SARS-CoV-2 has recently been evaluated in a phase 2b/3 efficacy trial in humans. CV2CoV is a second-generation mRNA vaccine with optimized non-coding regions and enhanced antigen expression. Here we report a head-to-head study of the immunogenicity and protective efficacy of CVnCoV and CV2CoV in nonhuman primates. We immunized 18 cynomolgus macaques with two doses of 12 ug of lipid nanoparticle formulated CVnCoV, CV2CoV, or sham (N=6/group). CV2CoV induced substantially higher binding and neutralizing antibodies, memory B cell responses, and T cell responses as compared with CVnCoV. CV2CoV also induced more potent neutralizing antibody responses against SARS-CoV-2 variants, including B.1.351 (beta), B.1.617.2 (delta), and C.37 (lambda). While CVnCoV provided partial protection against SARS-CoV-2 challenge, CV2CoV afforded robust protection with markedly lower viral loads in the upper and lower respiratory tract. Antibody responses correlated with protective efficacy. These data demonstrate that optimization of non-coding regions can greatly improve the immunogenicity and protective efficacy of an mRNA SARS-CoV-2 vaccine in nonhuman primates.

---

The CVnCoV mRNA vaccine (CureVac) has recently reported efficacy results in humans in the Phase 2b/3 HERALD trial in a population that included multiple viral variants. The observed vaccine efficacy against symptomatic COVID-19 was approximately 48% and 53% in the overall study population and in the 18-60 years of age subgroup, respectively [1]. CV2CoV is a second-generation mRNA vaccine that involves modifications of the non-coding regions that were selected based on an empiric screen for improved antigen expression [2, 3]. Both CVnCoV and CV2CoV are based on RNActive® technology [4–7] that consists of non-chemically modified, sequence engineered mRNA, without pseudouridine [6–12]. Both vaccines encode for the same full-length, pre-fusion stabilized SARS-CoV-2 Spike (S) [13, 14] and are encapsulated in lipid nanoparticles (LNP) with identical composition. CV2CoV has been engineered with different non-coding regions flanking the open reading frame, which have previously been shown to improve transgene expression [3] and protection against SARS-CoV-2 in ACE2 transgenic mice [2]. Specifically, CV2CoV includes 5’ UTR HSD17B4 and 3’ UTR PSMB3 elements, followed by a histone stem loop motif and a poly-A sequence (Fig. 1a; see Methods). In this study, we compare head-to-head the immunogenicity and protective efficacy of CVnCoV and CV2CoV against SARS-CoV-2 challenge in nonhuman primates.

**Figure 1.**
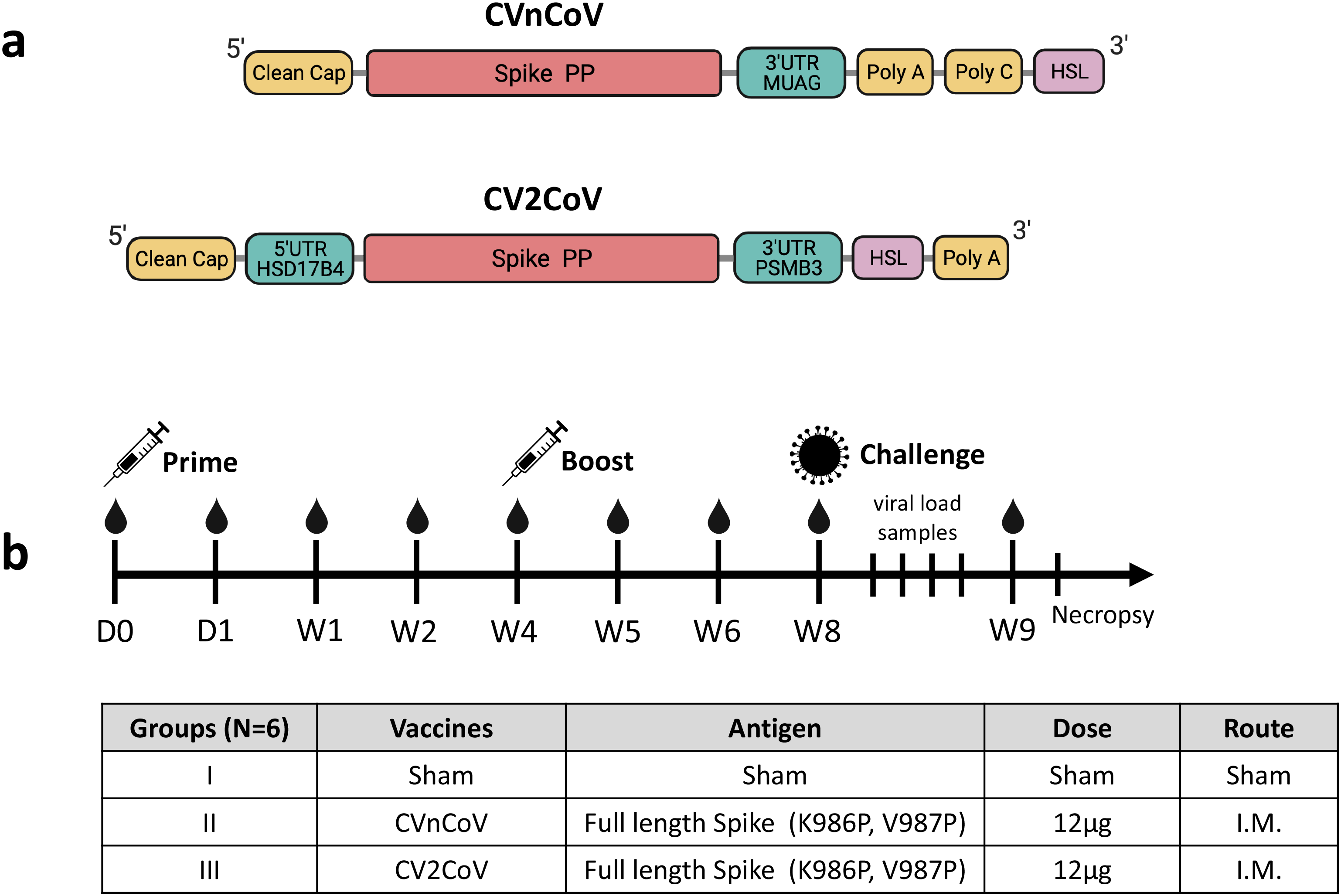
(a) mRNA vaccine design and (b) NHP vaccine study schema.

## Vaccine Immunogenicity

We immunized 18 cynomolgus macaques intramuscularly with either 12 µg CVnCoV or CV2CoV or sham vaccine (Fig. 1b). Animals were primed at week 0 and boosted at week 4. Sera was isolated from all animals 24h after the first vaccination to assess innate cytokine responses. CV2CoV induced higher levels of IFNɑ2a, IP-10 and MIP-1 compared with CVnCoV (P = 0.0152, P = 0.0152, P = 0.0411, respectively; Extended Data Fig. 1).

Binding antibody responses were assessed by receptor binding domain (RBD)-specific ELISAs at multiple timepoints following immunization [15, 16]. At week 2, binding antibody titers were only detected with CV2CoV and not with CVnCoV (CVnCoV median titer 25 [range 25-25]; CV2CoV median titer 799 [range 82-2,010]) (Fig. 2a). One week following the week 4 boost, antibody titers increased in both groups (CVnCoV median titer 48 [range 75-710]); CV2CoV median titer 28,407 [range 2,714-86,541]) (Fig. 2a). By week 8, binding antibody titers increased in the CVnCoV group but were still >50-fold lower than in the CV2CoV group (P=0.0043) (CVnCoV median titer 214 [range 47-1,238]; CV2CoV median titer 14,827 [range 2,133-37,079]).

**Figure 2.**
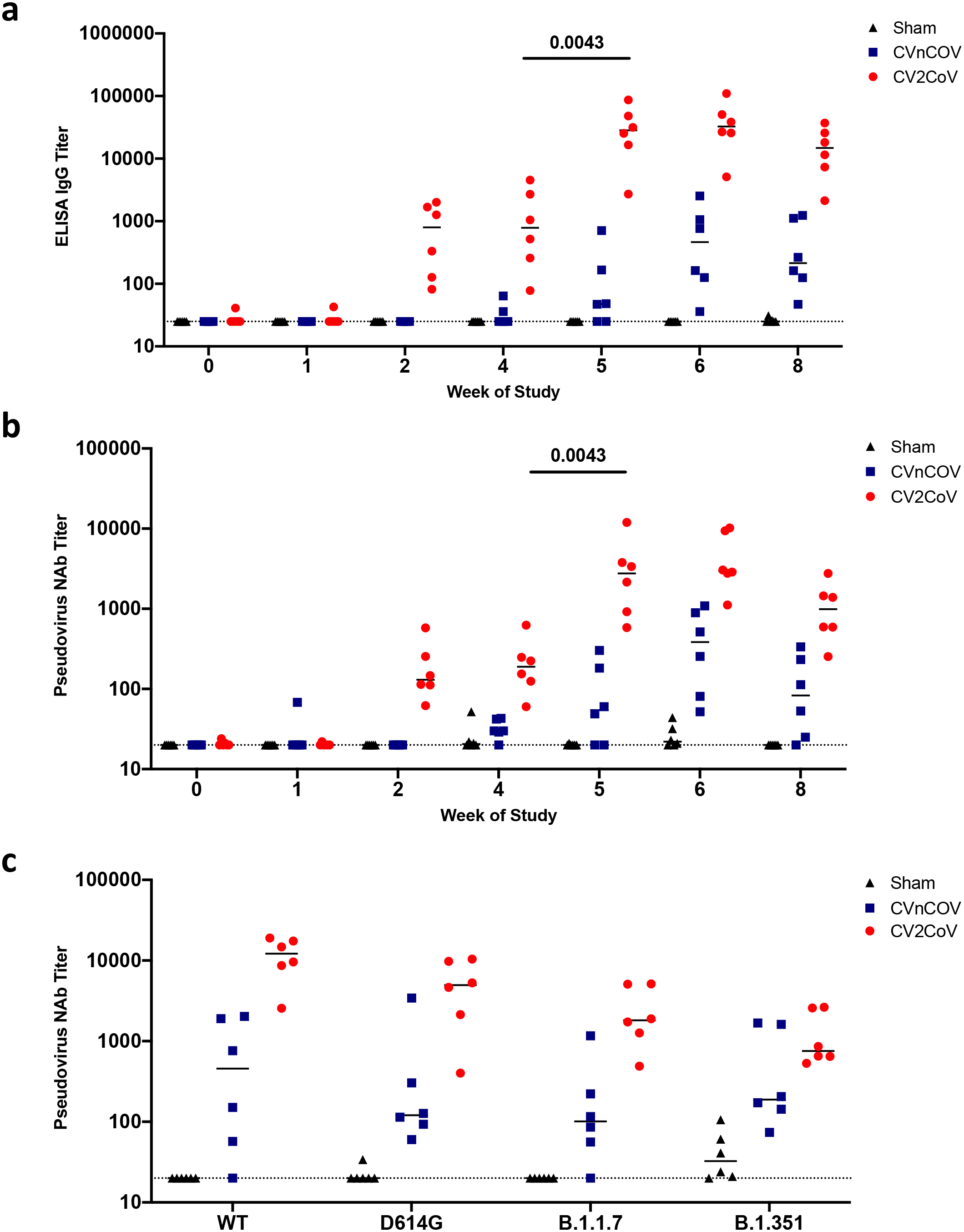
CV2CoV elicits high levels of binding and neutralizing antibody responses in NHPs. NHPs (6/group) were vaccinated twice with 12µg of CVnCoV or CV2CoV on d0 and d28 or remained untreated as negative controls (sham). (**a**) Titers of RBD binding antibodies and (**b**) pseudovirus neutralizing antibodies against ancestral SARS-CoV-2 strain were evaluated at different time points post first (week 0, 1, 2 and 4) and second (week 5, 6 and 8) vaccinations. (**c**) Sera isolated on d42 (week 6) were analyzed for pseudovirus neutralizing antibodies titers against the ancestral WA/2020 (WT) strain, virus featuring the D614G mutation, and variants including B.1.1.7 (Alpha) and B.1.351 (Beta). Each dot represents an individual animal, bars depict the median and the dotted line shows limit of detection.

Neutralizing antibody (NAb) responses were assessed by pseudovirus neutralization assays initially using the vaccine-matched SARS-CoV-2 wildtype (WT) WA1/2020 strain [15–17]. NAb titers followed a similar trend as binding antibody titers (Fig. 2b). At week 2, NAb were only detected with CV2CoV and not with CVnCoV (CVnCoV median titer 20 [range 20-20]; CV2CoV median titer 131 [range 62-578]) (Fig. 2b). One week following the week 4 boost, NAb titers increased (CVnCoV median titer 55 [range 20-302]; CV2CoV median titer 2,758 [range 583-11,941]). By week 8, NAb titers increased in the CVnCoV group but were still >10-fold lower than in the CV2CoV group (P=0.0043) (CVnCoV median titer 83 [range 20-335]; CV2CoV median titer 991 [range 253-2,765]).

At week 6, median NAb titers against the WT WA1/2020, D614G, B.1.1.7 (alpha), and B.1.351 (beta) variants were 456, 121, 101, and 189, respectively, for CVnCoV and were 12,181, 4962, 1813, and 755, respectively, for CV2CoV (Fig. 2c). Median NAb titers against C.37 (lamda), B.1.617.1 (kappa), and B.1.617.2 (delta) were 516, 158, and 36, respectively, for CVnCoV and were 1195, 541, and 568, respectively, for CV2CoV (Extended Data Fig. 2). Taken together, these data show that CV2CoV induced substantially higher NAb titers as well against SARS-CoV-2 variants compared with CVnCoV.

Most SARS-CoV-2 RBD-specific B cells reside within the memory B cell pool [18]. We assessed memory B cell responses in blood from CVnCoV, CV2CoV and sham vaccinated NHPs by flow cytometry [19]. RBD- and Spike-specific memory B cells were detected in the CV2CoV group, but not in the CVnCoV group at week 6 (Fig. 3a, 3b). The cells were not detected at week 1 for both groups (data not shown). T cell responses were assessed by IFN-γ enzyme-linked immunosorbent spot (ELISPOT) assay using pooled S peptides at week 6 in both groups but were higher in the CV2CoV group (P=0.0065) (Fig. 3c).

**Figure 3.**
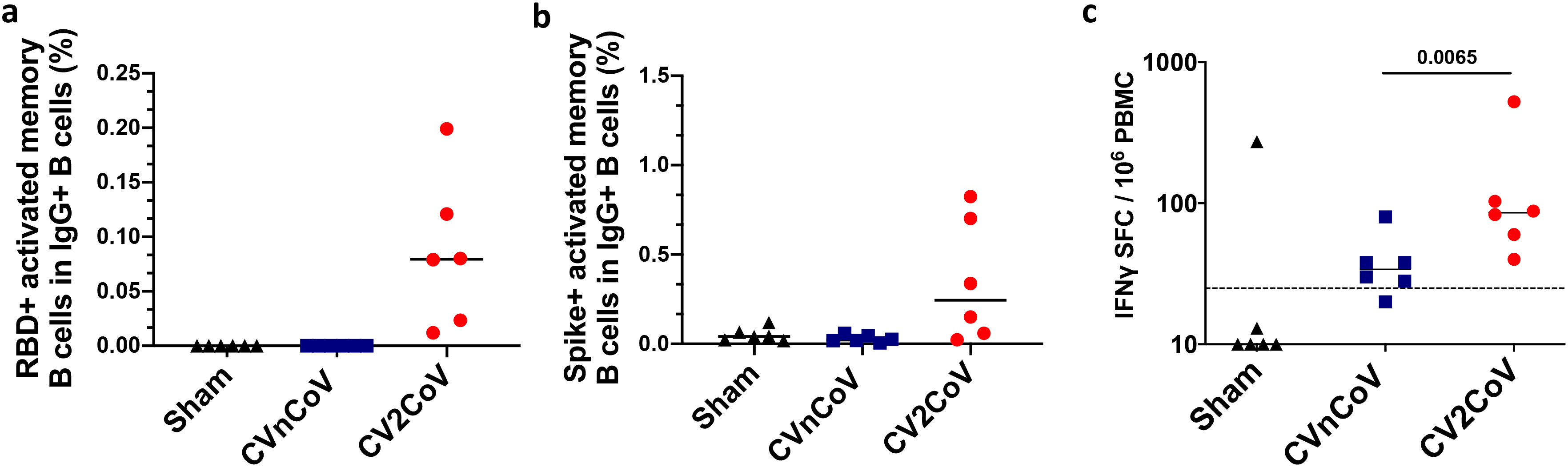
CV2CoV induces memory B cell and T cell immune responses in on day 42. PBMCs from negative control (sham), CVnCoV or CV2CoV vaccinated animals isolated on d42 of the experiment were stained for (**a**) RBD and (**b**) Spike-specific activated memory B cells and analyzed by high-parameter flow cytometry. IFNγ responses to pooled spike peptides were analyzed via ELISPOT (**c**). Each dot represents an individual animal, bars depict the median and the dotted line shows limit of detection. PBMC = peripheral blood mononuclear cell; SFC = spot forming cells

## Protective Efficacy

All animals were challenged at week 8 with 1.0×10^5^ TCID_50_ SARS-CoV-2 WA1/2020 via the intranasal (IN) and intratracheal (IT) routes. Viral loads were assessed in bronchoalveolar lavage (BAL) and nasal swab (NS) samples collected on days 1, 2, 4, 7 and 10 following challenge by RT-PCR specific for subgenomic mRNA (sgRNA) [20]. High subgenomic RNA levels were observed in BAL and NS in the sham group peak on day 2 and largely resolved by day 10. Sham controls had a peak median of 6.02 (range 4.62–6.81) log_10_ sgRNA copies/ml in BAL and 7.35 (range 5.84–8.09) log_10_ sgRNA copies/swab in NS on day 2 (Fig. 4). CVnCoV immunized animals showed a peak median of 4.92 (range 2.40–6.61) log_10_ sgRNA copies/ml in BAL and 6.42 (range 4.46–7.81) log_10_ sgRNA copies/swab in NS (Fig. 4). CV2CoV immunized animals exhibited a peak median of 2.90 (range 1.70–4.64) log_10_ sgRNA copies/ml in BAL and 3.17 (range 2.59–5.63) log_10_ sgRNA copies/swab in NS (Fig. 4), with resolution of sgRNA in BAL by day 2 in most animals and by day 4 in all animals. Overall, CV2CoV resulted in significantly lower peak viral loads than CVnCoV in both BAL (P=0.0411) and NS (P=0.0087) (Fig. 5a and b).

**Figure 4.**
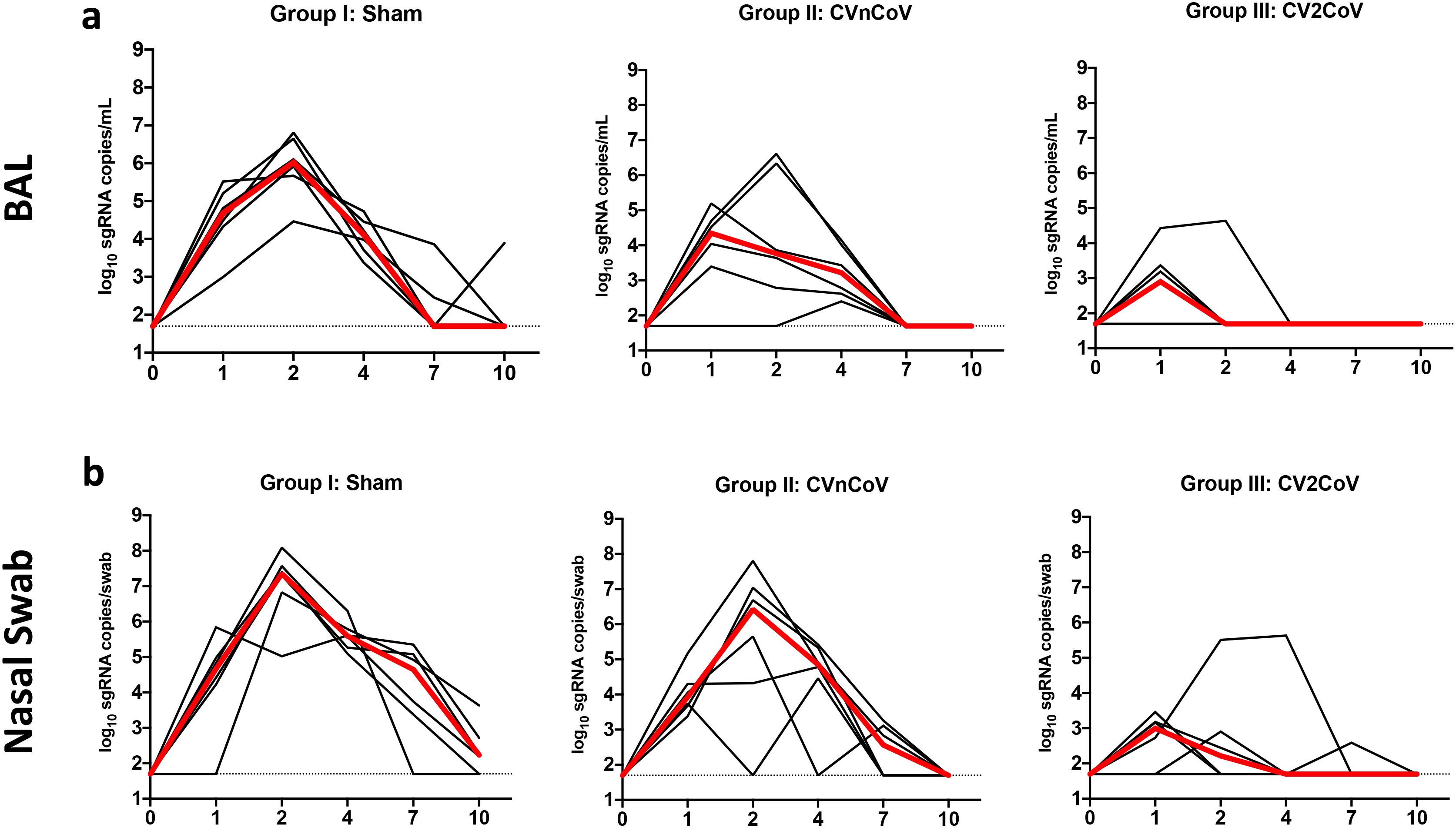
CV2CoV protects NHPs from challenge infection. Negative control (sham) or animals vaccinated on d0 and d28 of the experiment with 12µg of CVnCoV or CV2CoV as indicated were subjected to challenge infection using 1.0×10^5^ TCID_50_ SARS-CoV-2 via intranasal (IN) and intratracheal (IT) routes. BAL (**a**) and nasal swab samples (**b**) collected on days 1, 2, 4, 7 and 10 post-challenge were analyzed for levels of replicating virus by RT-PCR specific for subgenomic mRNA (sgRNA). Thin black lines represent an individual animal, thick red lines depict the median and the dotted line shows limit of detection. BAL = bronchoalveolar lavage

**Figure 5.**
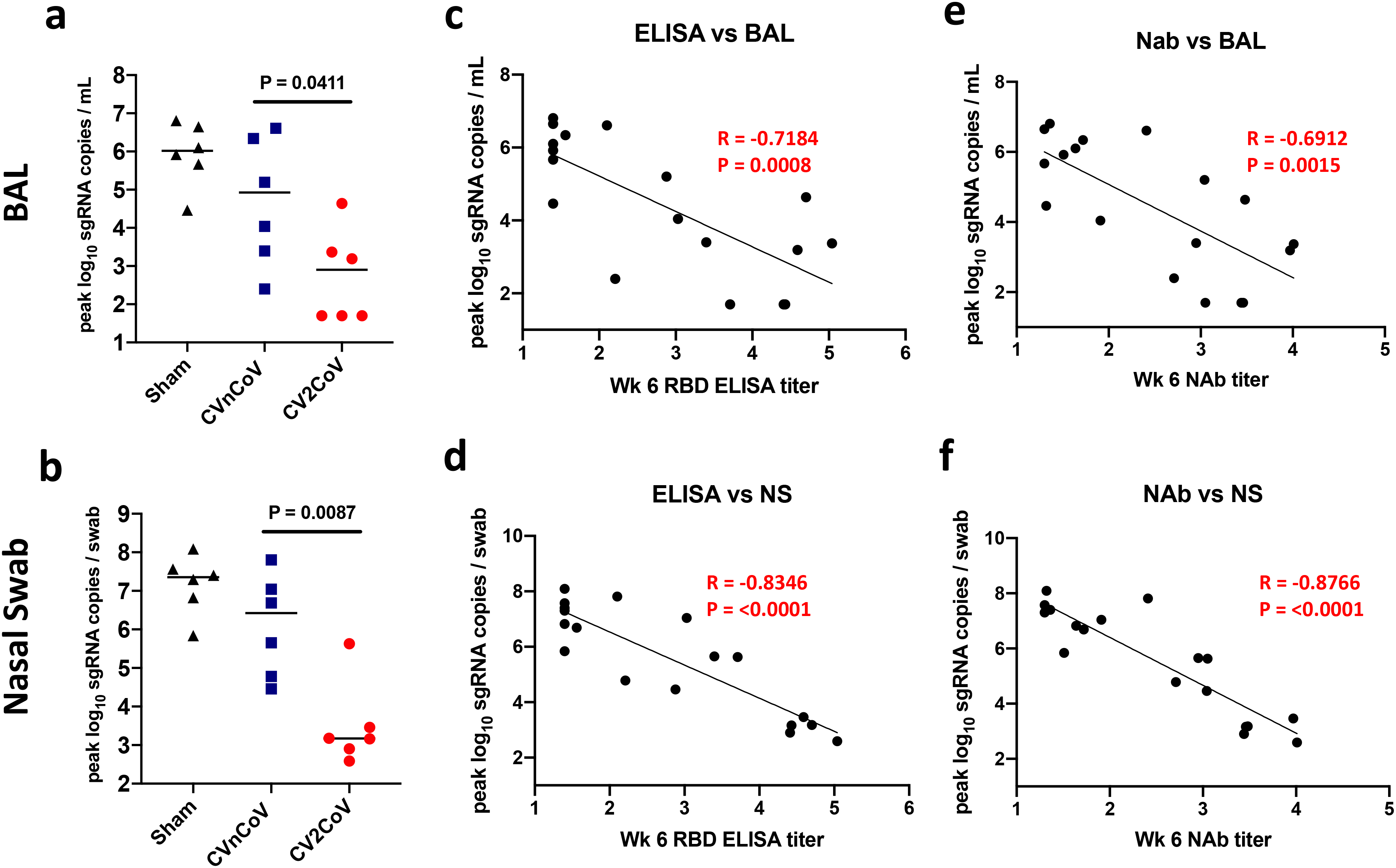
Titers of binding and neutralizing antibody titers elicited upon CVnCoV and CV2CoV vaccination correlate with protection against SARS-CoV-2. Summary of peak viral loads following SARS-CoV-2 challenge in BAL and Nasal Swab (left panels **a** and **b**); antibody correlates of protection for binding antibodies (middle panels in **c** and **d**) and neutralizing antibodies (right panels **e** and **f**). NAbs = neutralizing antibodies, BAL = bronchoalveolar lavage NS = nasal swab

We next evaluated immune correlates of protection in this study. The log_10_ ELISA and NAb titers at week 6 inversely correlated with peak log_10_ sgRNA copies/ml in BAL (P=0.0008, R=−0.7148 and P=0.0015, R= −0.6912, respectively, two-sided Spearman rank-correlation test) (Fig. 5, c and e) and with peak sgRNA copies/nasal swab in NS (P<0.0001, R=−0.8346, and P<0.0001, R=−0.8766, respectively, two-sided Spearman rank-correlation test) (Fig. 5, d and f). Consistent with prior observations from our laboratory and others [15, 16, 21], these findings suggest that binding and neutralizing antibody titers are important correlates of protection for these SARS-CoV-2 vaccines in nonhuman primates. Similar correlates of protection were observed with viral loads assessed as area under the curve (AUC) (Extended Data Fig. 3).

Following challenge, we observed anamnestic binding and neutralizing antibody responses in the CVnCoV vaccinated animals (Extended Data Fig. 4). We did not observe higher antibody responses in the CV2CoV vaccinated animals following challenge, likely reflecting the robust protection and minimal viral replication in these animals, as we have previously reported [16].

On day 10 post-challenge, animals were necropsied, and lung tissues were evaluated by histopathology. Although viral replication had largely resolved by this timepoint, sham animals had higher cumulative lung pathology scores [19] as compared to both CVnCoV and CV2CoV vaccinated animals (CVnCoV P= 0.0368; CV2CoV P= 0.0022) (Extended Data Fig. 5a). Sham animals also had more lung lobes affected (Extended Data Fig. 5b) and more extensive lung lesions with a greater proportion of lung lobes showing evidence of interstitial inflammation, alveolar inflammatory infiltrates, and type II pneumocyte hyperplasia (Extended Data Fig. 5c-h). No significant differences were observed between the cumulative lung scores between CVnCoV and CV2CoV vaccinated animals on day 10. Pathologic lesions in vaccinated animals were similar to those observed in sham animals (Extended Data Fig. 5i-l) but fewer overall and more focal in distribution.

## Discussion

CV2CoV elicited substantially higher humoral and cellular immune responses and provided significantly improved protective efficacy against SARS-CoV-2 challenge compared with CVnCoV in macaques. These data suggest that optimization of non-coding elements of the mRNA backbone can substantially improve the immunogenicity and protective efficacy of mRNA vaccines. Both CVnCoV and CV2CoV contain non-modified nucleotides, without pseudouridine or derivates, and CV2CoV has previously been shown to lead to higher antigen expression than CVnCoV in cell culture [3]. While previous studies in rodents and nonhuman primates have demonstrated protection by CVnCoV [2, 22][23] this was only studied in the lower respiratory tract [22][23]. In the present study, CVnCoV provided only modest reductions in viral loads in BAL and NS compared with sham controls. In contrast, CV2CoV induced >10-fold higher NAb responses than CVnCoV against multiple viral variants and provided >3 log reductions in sgRNA copies/ml in BAL and >4 log reductions in sgRNA copies/swab in NS compared with sham controls.

Previous mRNA vaccine clinical trials have demonstrated onset of protective efficacy after the first dose and improved protection after the boost immunization [24, 25]. In the present study, the prime immunization with CV2CoV induced binding and neutralizing antibodies in all macaques by week 2, and these responses increased substantially by 1 week after the boost immunization. Although comparisons with other nonhuman primate studies are difficult due to differences in study designs and laboratory assays, NAb titers induced by CV2CoV appear roughly similar to those reported for previous mRNA vaccines in macaques [26, 27].

As previously reported for other vaccines [28–32], NAb titers were lower to certain SARS-CoV-2 variants, including B.1.351 (beta) and B.1.617.2 (delta), than to the parental strain WA1/2020. Although our challenge virus in this study was SARS-CoV-2 WA1/2020, NAb titers elicited by CV2CoV to these viral variants exceeded the threshold that we previously reported as threshold titers for protection (50-100) [17, 19, 21]. However, future studies will be required to assess directly the protective efficacy of CV2CoV against SARS-CoV-2 variants of concern in non-human primates.

CV2CoV induced both antigen-specific memory B cell responses and T cell responses. While the correlates of protection in this study were binding and neutralizing antibodies [33, 34], it is likely that CD8^+^ T cells contribute to viral clearance in tissues [35, 36]. We previously reported that depletion of CD8^+^ T cells partially abrogated protective efficacy against SARS-CoV-2 re-challenge in convalescent macaques [21]. Memory B cells may contribute to durability of antibody responses [37, 38], although B cell germinal center responses and durability of protective efficacy following CV2CoV vaccination remain to be determined. Moreover, although this study was not specifically designed as a safety study, it is worth noting that we did not observe any adverse effects following CVnCoV or CV2CoV vaccination, and we did not observe any unexpected or enhanced pathology in the vaccinated animals at necropsy [39].

In summary, our data show that optimization of non-coding regions in a SARS-CoV-2 mRNA vaccine can substantially improve its immunogenicity against multiple viral variants and enhance protective efficacy against SARS-CoV-2 challenge in nonhuman primates. Improved characteristics of CV2CoV, compared with CVnCoV, could translate into increased efficacy in humans, and clinical trials of CV2CoV are planned.

## Data availability statement

All data are available in the manuscript and the supplementary material. This work is licensed under a Creative Commons Attribution 4.0 International (CC BY 4.0) license, which permits unrestricted use, distribution, and reproduction in any medium, provided the original work is properly cited. To view a copy of this license, visit https://creativecommons.org/licenses/by/4.0/. This license does not apply to figures/photos/artwork or other content included in the article that is credited to a third party; obtain authorization from the rights holder before using such material.

## Acknowledgements

We thank Sarah Gardner, Gabriella Kennedy, and Rachael Edmonston for their generous assistance. We thank Domenico Maione and Marie-Thérèse Martin for critically reading the manuscript.

## Author contributions

S.R., B.P., S.O.M., N.R. and D.H.B. designed the study. M.S.G., J.Y., A.C., N. M., X. H., J. L., K. M., A. M., T.G., D.H., S.P., D.S., O.S., and J.B. performed immunologic and virologic assays. X.L. and A.C.C. performed cytokine analysis. L.P., D.V., Z.F., J.Y., J.M., R.B., A.C., E.T., H.A. and M.L. led the clinical care of the animals. M.S.G. and D.H.B. wrote the paper with all coauthors.

## Corresponding authors

Dan H. Barouch, M.D.; Susanne Rauch, Ph.D.

## Materials and Methods

### mRNA vaccines

The two mRNA vaccines, CVnCoV and CV2CoV, are based on CureVac’s RNActive® platform (claimed and described in e.g. WO2002098443 and WO2012019780) and do not include chemically modified nucleosides. They are comprised of a 5ʹ cap1 structure, a GC-enriched open reading frame (ORF), 3ʹ UTR and a vector-encoded poly-A stretch. CVnCoV contains a cleanCap (Trilink), parts of the 3’ UTR of the Homo sapiens alpha haemoglobin gene as 3’ UTR, followed by a poly-A (64) stretch, a polyC (30) stretch and a histone stem loop [22, 23]. CV2CoV has previously been described to contain a cleanCap followed by 5’ UTR from the human hydroxysteroid 17-beta dehydrogenase 4 gene (HSD17B4) and a 3’ UTR from human proteasome 20S subunit beta 3 gene (PSMB3), followed by a histone stem loop and a poly-A (100) stretch [3]. Both constructs were encapsulated in lipid nanoparticles (LNP) by Acuitas Therapeutics (Vancouver, Canada) (CV2CoV) or Polymun Scientific Immunbiologische Forschung GmbH (Klosterneuburg, Austria) (CVnCoV). LNPs are composed of ionizable amino lipid, phospholipid, cholesterol, and a PEGylated lipid; compositions for CVnCoV and CV2CoV are identical. Both mRNAs encode for SARS-CoV-2 full length spike protein containing stabilizing K986P and V987P mutations (NCBI Reference Sequence NC_045512.2).

### Animals and study design

18 cynomolgus macaques were randomly assigned to three groups. Animals received either CVnCoV (N=6) or CV2CoV (N=6) mRNA vaccines or were designated as sham controls (N=6). The mRNA vaccines were administered at a 12 µg dose, intramuscularly, in the left quadriceps on day 0. Boost immunizations were similarly administered at week 4. At week 8, all animals were challenged with 1.0×10^5^ TCID_50_ SARS-CoV-2 derived from USA-WA1/2020 (NR-52281; BEI Resources) [17]. Challenge virus was administered as 1 ml by the intranasal (IN) route (0.5 ml in each nare) and 1 ml by the intratracheal (IT) route. All animals were sacrificed 10 days post challenge. Immunologic and virologic assays were performed blinded. All animals were housed at Bioqual, Inc. (Rockville, MD). All animal studies were conducted in compliance with all relevant local, state, and federal regulations and were approved by the Bioqual Institutional Animal Care and Use Committee (IACUC).

### Cytokine analyses

Serum levels of 19 analytes that have been associated with immune response to viral infection were tested using U-PLEX Viral Combo 1 (NHP) kit (K15069L-1) from Meso Scale Discovery (MSD, Rockville, MD). The 19 analytes and their detection limits (LLODs) are G-CSF (1.5 pg/mL), GM-CSF (0.12 pg/mL), IFN-α2a (1.7 pg/mL), IFN-γ (1.7 pg/mL), IL-1RA (1.7 pg/mL), IL-1β (0.15 pg/mL), IL-4 (0.06 pg/mL), IL-5 (0.24 pg/mL), IL-6 (0.33 pg/mL), IL-7 (1.5 pg/mL) and IL-8 (0.15 pg/mL), IL-9 (0.14 pg/mL), IL-10 (0.14 pg/mL), IL-12p70 (0.54 pg/mL), IP-10 (0.49 pg/mL), MCP-1 (0.74 pg/mL), MIP-1α (7.7 pg/mL), TNF-α (0.54 pg/mL) and VEGF-A (2.0pg/mL). All serum samples were assayed in duplicate. Assay was done by the Metabolism and Mitochondrial Research Core (Beth Israel Deaconess Medical Center, Boston, MA) following manufacture’s instruction. The assay plates were read by MESO QUICKPLEX SQ 120 instrument and data were analyzed by DISCOVERY WORKBENCH® 4.0 software.

### Enzyme-linked immunosorbent assay (ELISA)

RBD-specific binding antibodies were assessed by ELISA as described [16, 17]. Briefly, 96-well plates were coated with 1μg/ml SARS-CoV-2 RBD protein (40592-VNAH, SinoBiological) in 1X DPBS and incubated at 4°C overnight. After incubation, plates were washed once with wash buffer (0.05% Tween 20 in 1 X DPBS) and blocked with 350 μL Casein block/well for 2–3 h at room temperature. After incubation, block solution was discarded, and plates were blotted dry. Serial dilutions of heat-inactivated serum diluted in casein block were added to wells and plates were incubated for 1 h at room temperature. Next, the plates were washed three times and incubated for 1 h with a 1:1000 dilution of anti-macaque IgG HRP (NIH NHP Reagent Program) at room temperature in the dark. Plates were then washed three more times, and 100 μL of SeraCare KPL TMB SureBlue Start solution was added to each well; plate development was halted by the addition of 100 μL SeraCare KPL TMB Stop solution per well. The absorbance at 450nm was recorded using a VersaMax or Omega microplate reader. ELISA endpoint titers were defined as the highest reciprocal serum dilution that yielded an absorbance > 0.2. Log10 endpoint titers are reported. Immunologic assays were performed blinded.

### Pseudovirus neutralization assay

The SARS-CoV-2 pseudoviruses expressing a luciferase reporter gene were generated as described previously [15, 40]. Briefly, the packaging plasmid psPAX2 (AIDS Resource and Reagent Program), luciferase reporter plasmid pLenti-CMV Puro-Luc (Addgene), and Spike protein expressing pcDNA3.1-SARS CoV-2 SΔCT of variants were co-transfected into HEK293T cells by lipofectamine 2000 (ThermoFisher). Pseudoviruses of SARS-CoV-2 variants were generated by using WA1/2020 strain (Wuhan/WIV04/2019, GISAID accession ID: EPI_ISL_402124), D614G mutation, B.1.1.7 variant (GISAID accession ID: EPI_ISL_601443), B.1.351 variant (GISAID accession ID: EPI_ISL_712096), C37 variant (GenBank ID: QRX62290), B.1.671.1 variant (GISAID accession ID: EPI_ISL_1384866) and B.1.617.2 variant (GISAID accession ID: EPI_ISL_2020950). The supernatants containing the pseudotype viruses were collected 48 h post-transfection, which were purified by centrifugation and filtration with 0.45 µm filter. To determine the neutralization activity of the plasma or serum samples from participants, HEK293T-hACE2 cells were seeded in 96-well tissue culture plates at a density of 1.75 x 10^4^ cells/well overnight. Three-fold serial dilutions of heat inactivated serum or plasma samples were prepared and mixed with 50 µL of pseudovirus. The mixture was incubated at 37°C for 1 h before adding to HEK293T-hACE2 cells. 48 h after infection, cells were lysed in Steady-Glo Luciferase Assay (Promega) according to the manufacturer’s instructions. SARS-CoV-2 neutralization titers were defined as the sample dilution at which a 50% reduction in relative light unit (RLU) was observed relative to the average of the virus control wells.

### B cell immunophenotyping

Fresh PBMCs were stained with Aqua live/dead dye (Invitrogen) for 20 min, washed with 2% FBS/DPBS buffer, and suspended in 2% FBS/DPBS buffer with Fc Block (BD) for 10 min, followed by staining with monoclonal antibodies against CD45 (clone D058-1283, BUV805), CD3 (clone SP34.2, APC-Cy7), CD7 (clone M-T701, Alexa700), CD123 (clone 6H6, Alexa700), CD11c (clone 3.9, Alexa700), CD20 (clone 2H7, PE-Cy5), IgA (goat polyclonal antibodies, APC), IgG (clone G18-145, BUV737), IgM (clone G20-127, BUV396), IgD (goat polyclonal antibodies, PE), CD80 (clone L307.4, BV786), CD95 (clone DX2, BV711), CD27 (clone M-T271, BUV563), CD21 (clone B-ly4, BV605), CD14 (clone M5E2, BV570) and CD138 (clone DL-101, PE-CF594). Cells were also stained with SARS-CoV-2 antigens including biotinylated SARS-CoV-2 RBD protein (Sino Biological) and full-length SARS-CoV-2 Spike protein (Sino Biological) labeled with FITC and DyLight 405 (DyLight® 405 Conjugation Kit, FITC Conjugation Kit, Abcam), at 4 °C for 30 min. After staining, cells were washed twice with 2% FBS/DPBS buffer, followed by incubation with BV650 streptavidin (BD Pharmingen) for 10min, then washed twice with 2% FBS/DPBS buffer. After staining, cells were washed and fixed by 2% paraformaldehyde. All data were acquired on a BD FACSymphony flow cytometer. Subsequent analyses were performed using FlowJo software (Treestar, v.9.9.6). Immunologic assays were performed blinded.

### IFN-γ enzyme-linked immunospot (ELISPOT) assay

ELISPOT plates were coated with mouse anti-human IFN-γ monoclonal antibody from BD Pharmingen at a concentration of 5 μg/well overnight at 4°C. Plates were washed with DPBS containing 0.25% Tween 20, and blocked with R10 media (RPMI with 11% FBS and 1.1% penicillin-streptomycin) for 1 h at 37°C. The Spike 1 and Spike 2 peptide pools (JPT Peptide Technologies, custom made) used in the assay contain 15 amino acid peptides overlapping by 11 amino acids that span the protein sequence and reflect the N- and C-terminal halves of the protein, respectively. Spike 1 and Spike 2 peptide pools were prepared at a concentration of 2 μg/well, and 200,000 cells/well were added. The peptides and cells were incubated for 18–24 h at 37°C. All steps following this incubation were performed at room temperature. The plates were washed with ELISPOT wash buffer and incubated for 2 h with Rabbit polyclonal anti-human IFN-γ Biotin from U-Cytech (1 μg/mL). The plates are washed a second time and incubated for 2 h with Streptavidin-alkaline phosphatase antibody from Southern Biotechnology (1 μg/mL). The final wash was followed by the addition of Nitro-blue Tetrazolium Chloride/5-bromo-4-chloro 3 ‘indolyl phosphate p-toludine salt (NBT/BCIP chromagen) substrate solution (Thermo Scientific) for 7 min. The chromagen was discarded and the plates were washed with water and dried in a dim place for 24 h. Plates were scanned and counted on a Cellular Technologies Limited Immunospot Analyzer.

### Subgenomic RT-PCR assay

SARS-CoV-2 E gene subgenomic RNA (sgRNA) was assessed by RT-PCR using primers and probes as previously described [15, 17]. A standard was generated by first synthesizing a gene fragment of the subgenomic E gene [41]. The gene fragment was subsequently cloned into a pcDNA3.1+ expression plasmid using restriction site cloning (Integrated DNA Technologies). The insert was *in vitro* transcribed to RNA using the AmpliCap-Max T7 High Yield Message Maker Kit (CellScript). Log dilutions of the standard were prepared for RT-PCR assays ranging from 1×10^10^ copies to 1×10^-1^ copies. Viral loads were quantified from bronchoalveolar lavage (BAL) fluid and nasal swabs (NS). RNA extraction was performed on a QIAcube HT using the IndiSpin QIAcube HT Pathogen Kit according to manufacturer’s specifications (Qiagen). The standard dilutions and extracted RNA samples were reverse transcribed using SuperScript VILO Master Mix (Invitrogen) following the cycling conditions described by the manufacturer. A Taqman custom gene expression assay (Thermo Fisher Scientific) was designed using the sequences targeting the E gene sgRNA [41]. The sequences for the custom assay were as follows, forward primer, sgLeadCoV2.Fwd: CGATCTCTTGTAGATCTGTTCTC, E_Sarbeco_R: ATATTGCAGCAGTACGCACACA, E_Sarbeco_P1 (probe): VIC-ACACTAGCCATCCTTACTGCGCTTCG-MGBNFQ. Reactions were carried out in duplicate for samples and standards on the QuantStudio 6 and 7 Flex Real-Time PCR Systems (Applied Biosystems) with the thermal cycling conditions: initial denaturation at 95°C for 20 seconds, then 45 cycles of 95°C for 1 second and 60°C for 20 seconds. Standard curves were used to calculate subgenomic RNA copies per ml or per swab. The quantitative assay sensitivity was determined as 50 copies per ml or per swab.

### Histopathology

At time of fixation, lungs were suffused with 10% formalin to expand the alveoli. All tissues were fixed in 10% formalin and blocks sectioned at 5 µm. Slides were baked for 30-60 min at 65 degrees, deparaffinized in xylene, rehydrated through a series of graded ethanol to distilled water, then stained with hematoxylin and eosin (H&E). Blinded histopathological evaluation was performed by a board-certified veterinary pathologist (AJM).

### Statistical analyses

Statistical analyses were performed using GraphPad Prism (version 9.0) software (GraphPad Software) and comparison between groups was performed using a two-tailed nonparametric Mann-Whitney U t test. P-values of less than 0.05 were considered significant. Correlations were assessed by two-sided Spearman rank-correlation tests.

## Extended Data Figure Legends

**Extended Data Figure. 1.**
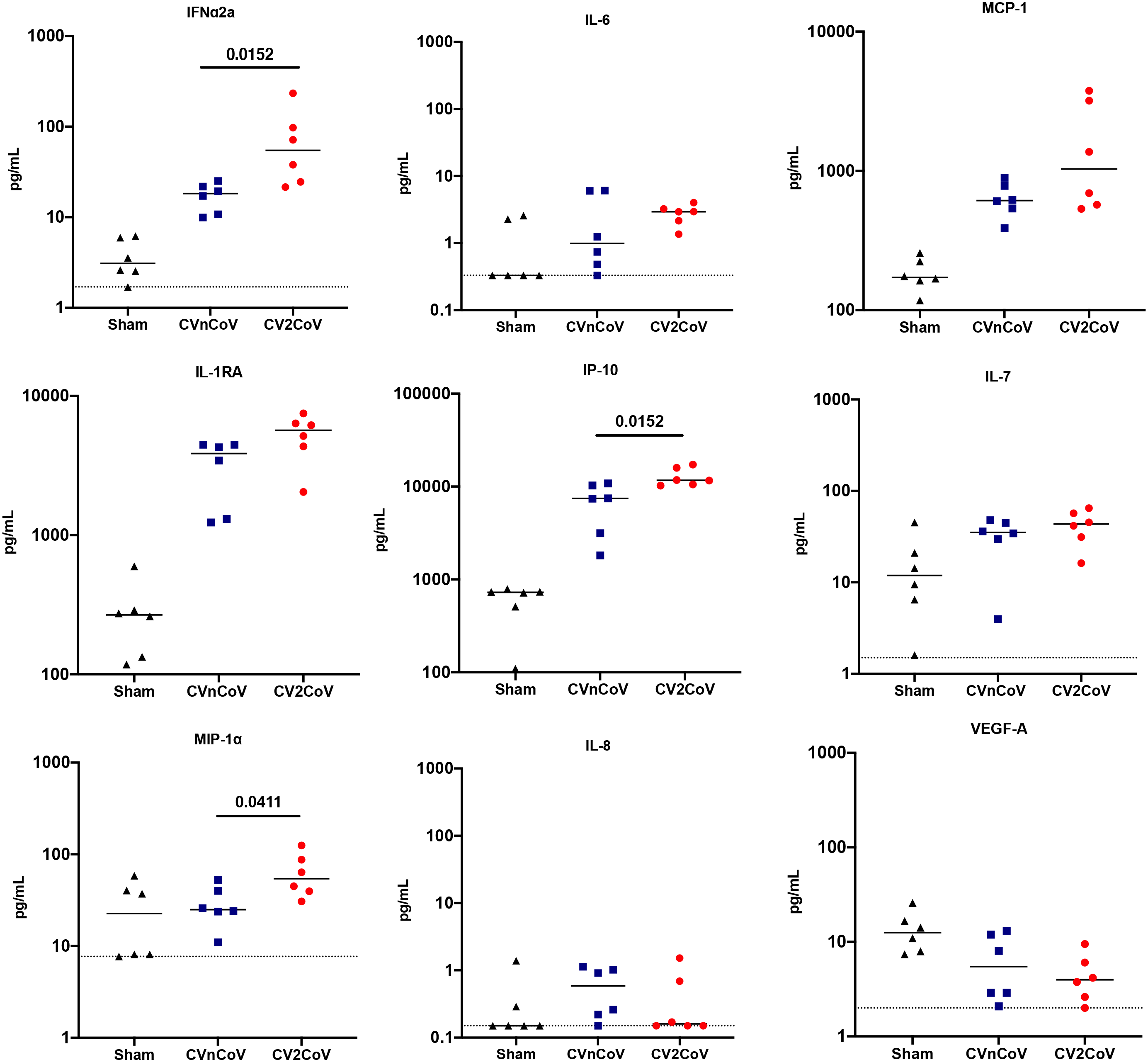
mRNA vaccination leads to innate cytokine induction in the serum of NHPs 24h post immunization. Sera isolated 24h post first injection were analyzed for a panel of 19 cytokines associated with viral infection using a U-PLEX Viral Combo kit from Meso Scale Discovery. Changes in cytokine levels above the detection limits were detectable for 9 cytokines. Each dot represents an individual animal, bars depict the median and the dotted line shows limit of detection. Statistical analysis was performed using Mann-Whitney test.

**Extended Data Figure. 2.**
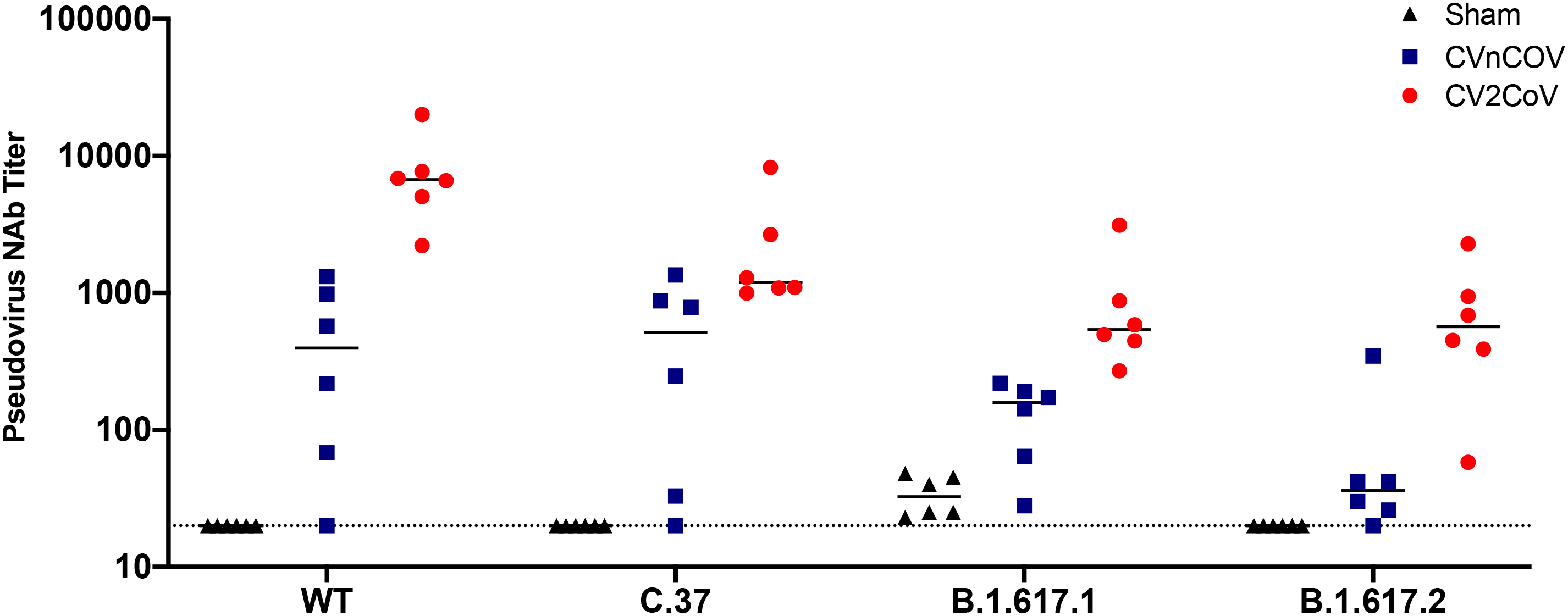
CV2CoV elicits high levels of binding and neutralizing antibody responses in NHPs. NHPs (6/group) were vaccinated twice with 12µg of CVnCoV or CV2CoV on d0 and d28 or remained untreated as negative controls (sham). Sera isolated on d42 (week 6) were analyzed for pseudovirus neutralizing antibodies titers against the ancestral WA/2020 (WT) strain, C.37 (Lambda), B.1.617.1 (Kappa) and B.1.617.2 (Delta). Each dot represents an individual animal, bars depict the median and the dotted line shows limit of detection.

**Extended Data Figure. 3.**
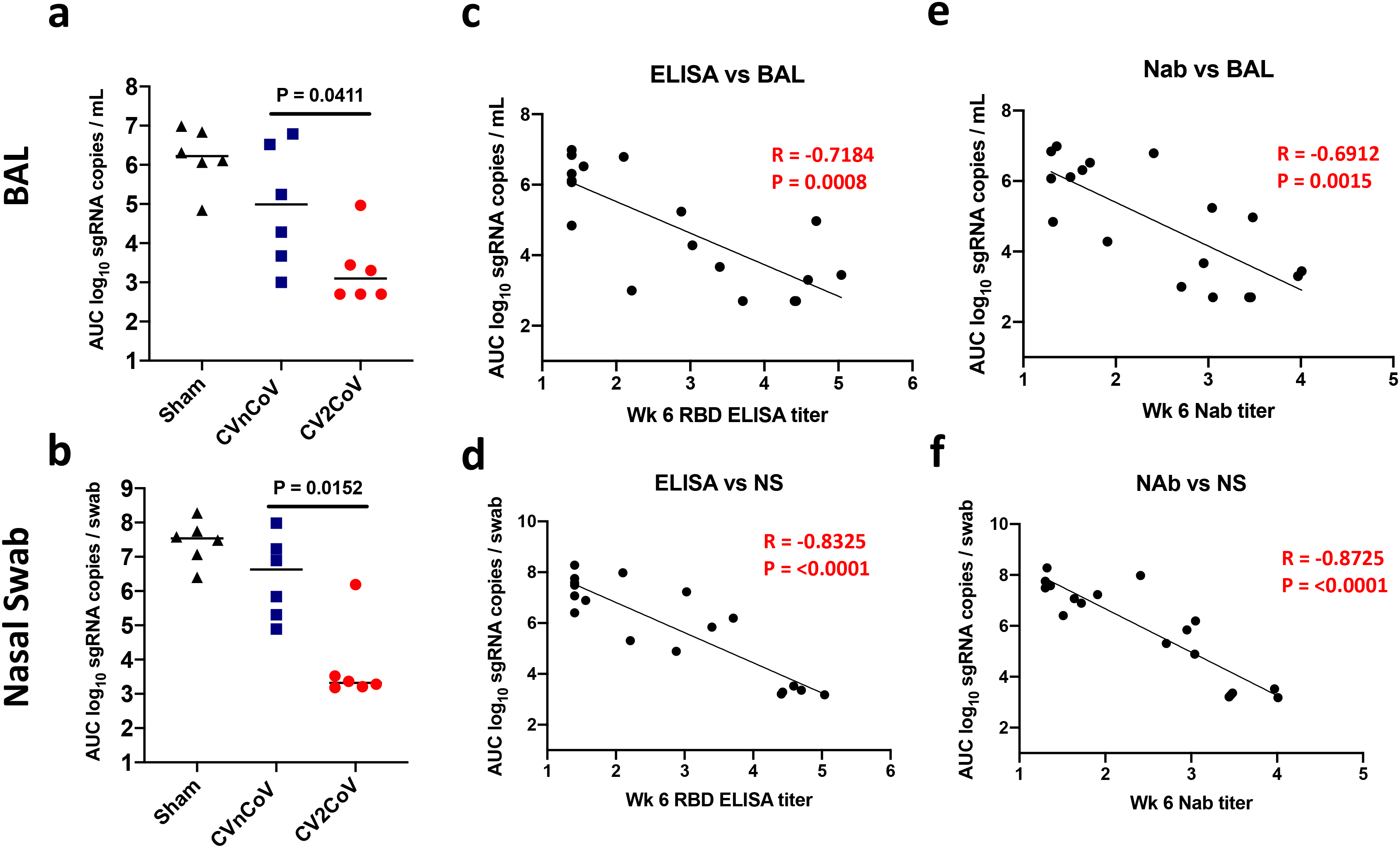
Titers of binding and neutralizing antibody titers elicited upon CVnCoV and CV2CoV vaccination correlate with protection against SARS-CoV-2. Summary of area under curve (AUC) viral load values following SARS-CoV-2 challenge in BAL and Nasal Swab (left panels **a** and **b**); antibody correlates of protection for binding antibodies (middle panels in **c** and **d**) and neutralizing antibodies (right panels **e** and **f**). NAbs = neutralizing antibodies, BAL = bronchoalveolar lavage NS = nasal swab

**Extended Data Figure. 4.**
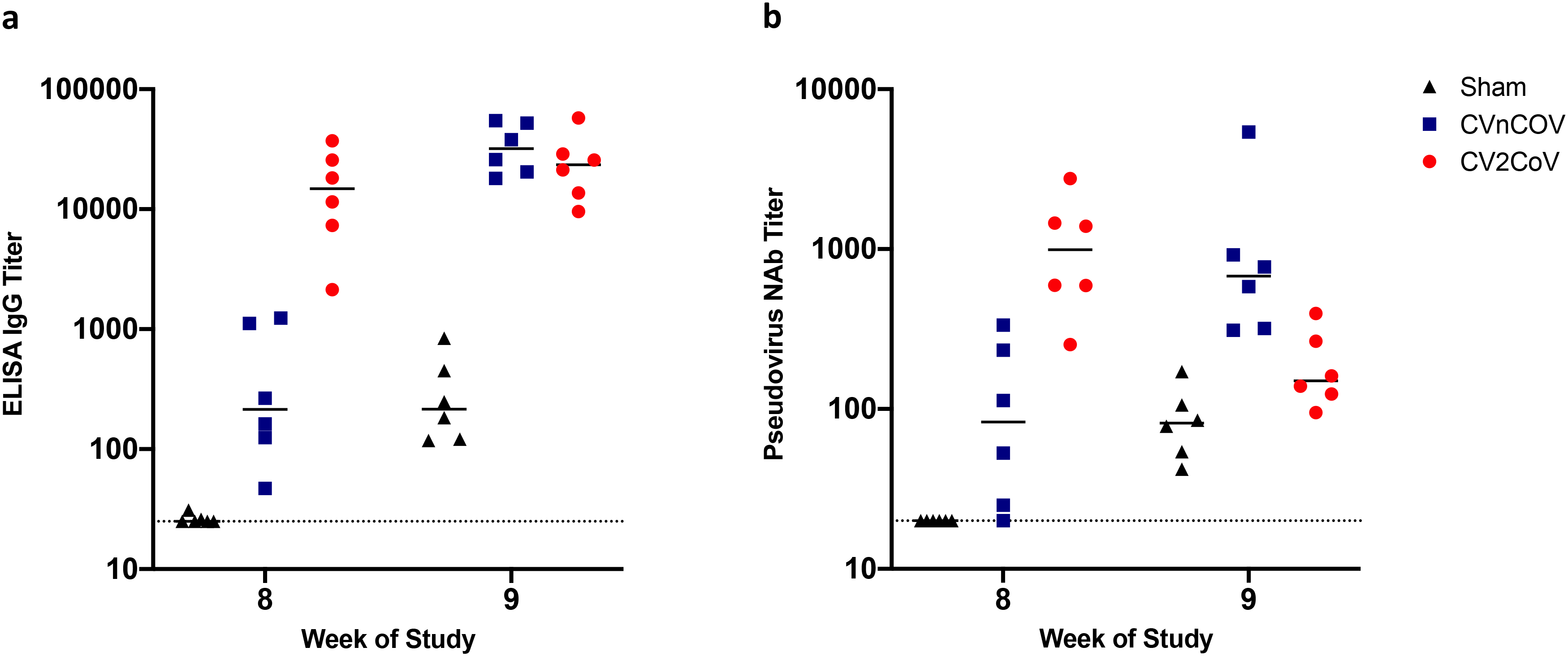
Post-challenge binding and neutralizing antibody responses of NHPs. Negative control (sham) or animals vaccinated on d0 and d28 of the experiment with 12µg of CVnCoV or CV2CoV as indicated were subjected to challenge infection using 1.0×10^5^ TCID_50_ SARS-CoV-2 via intranasal (IN) and intratracheal (IT) routes.). (**a**) Titers of RBD binding antibodies and (**b**) pseudovirus neutralizing antibodies against ancestral SARS-CoV-2 strain were evaluated before (week 8) and a week after challenge infection (week 9). Each dot represents an individual animal, bars depict the median and the dotted line shows limit of detection. NAbs= neutralizing antibodies

**Extended Data Figure. 5:**
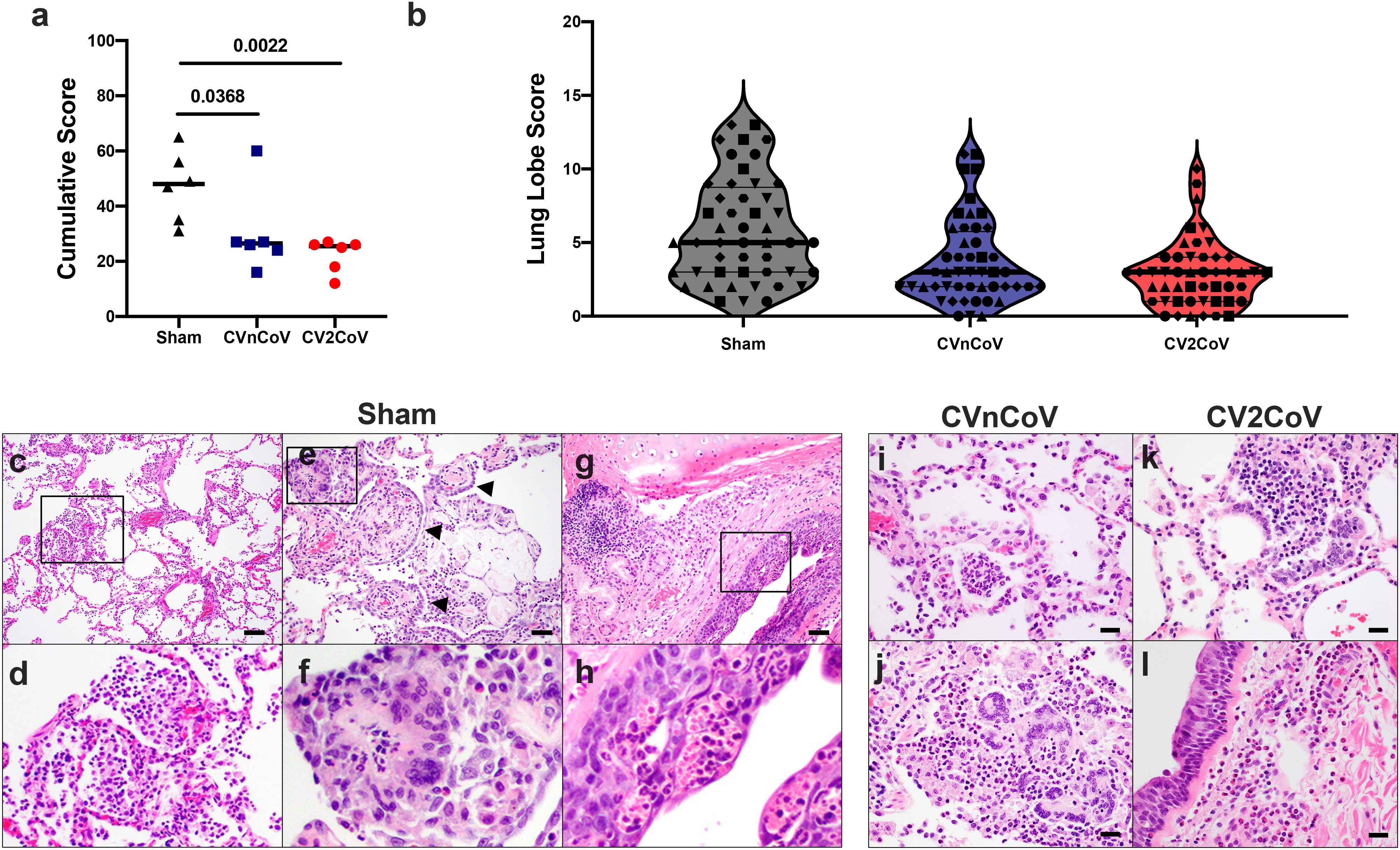
CVnCoV and CV2CoV protect the lungs from pathological changes upon viral challenge. Eight lung lobes (4 sections from right and left, caudal to cranial) were assessed and scored (1-4) for each of the following lesions: 1) Interstitial inflammation and septal thickening 2) Eosinophilic interstitial infiltrate 3) Neutrophilic interstitial infiltrate 4) Hyaline membranes 5) Interstitial fibrosis 6) Alveolar infiltrate, macrophage 7) Alveolar/Bronchoalveolar infiltrate, neutrophils 8) Syncytial cells 9) Type II pneumocyte hyperplasia 10) Broncholar infiltrate, macrophage 11) Broncholar infiltrate, neutrophils 12) BALT hyperplasia 13) Bronchiolar/peribronchiolar inflammation 14) Perivascular, mononuclear infiltrates 15) Vessels, endothelialitis. Each feature assessed was assigned a score of 0= no significant findings; 1=minimal; 2= mild; 3=moderate; 4=marked/severe. (**a**) Cumulative scores per animal (**b**) Cumulative scores per lung lobe. Individual animals are represented by symbols. Representative histopathology from sham vaccinated (**c-h**), CnVCoV vaccinated (**i, j**), and Cv2CoV vaccinated (**k, l**) animals showing (**c, d,** inset) alveolar macrophage infiltrate, (**e, f,** inset) syncytial cells (arrowheads) and type II pneumocyte hyperplasia, inset (**g, h,** inset) bronchiolar epithelial necrosis with neutrophilic infiltrates (**i**) alveolar neutrophilic infiltrate and alveolar septal thickening (**j**) focal consolidation with inflammation composed of macrophages, neutrophils, and syncytial cells (**k**) focal pneumocyte hyperplasia, syncytial cells and inflammatory infiltrates (**l**) peribronchiolar inflammation. Scale bars: 100 microns (**c**), 50 microns (**e, g**) 20 microns (**i-l**). BALT bronchus associated lymphoid tissue.

